# Low temperature abolishes human cellular circadian rhythm through Hopf bifurcation

**DOI:** 10.1101/2025.03.21.644557

**Authors:** Yaoyao Xiao, Yuko Sainoo, Takayuki Nishimura, Hiroshi Ito

## Abstract

Circadian clocks orchestrate behavior, physiology, and metabolism in harmony with the Earth’s 24-h cycle. Low temperatures are known to disrupt circadian clocks in plants and poikilotherms; however, their effects on human circadian rhythms remain poorly understood. Here, we demonstrate that cold exposure abolishes the circadian rhythm in cultured human cells through diminishing the oscillation amplitude, which was restored upon rewarming. In addition, the oscillation amplitude of the 24-h temperature cycles was enhanced through resonance, reflecting the intrinsic frequency of the circadian clock. From a theoretical perspective, these dynamics correspond to Hopf bifurcation, which is confirmed by a mathematical model for the mammalian circadian clock. In contrast, the circadian amplitude of human hair follicle cells was resistant to temperature changes, indicating robust temperature homeostasis. These findings suggest that Hopf bifurcation in the circadian clock for temperature regulation is shared by homeothermic and poik-ilothermic species.

## INTRODUCTION

The circadian rhythm is an endogenous oscillation that exists in almost all organisms. It has a period of approximately 24 h to adapt to the earth’s rotational period.^1^ Temperature is a major environmental factor that regulates circadian rhythms.^2^ Among circadian clock systems, temperature compensation and entrainment to environmental temperature cycles are prominent properties with distinct molecular mechanisms. However, the effect of low temperature exposure on circadian rhythms remains underexplored. Some reports in plants and poikilothermic animals have documented the nullification of circadian rhythms under low temperatures. For example, the circadian rhythm was abolished in the dinoflagellate *Lingulodinium polyedra* at 11.5 °C,^3^ in the duckweed *Lemna gibba* at 12 °C,^4^ in *Drosophila* at 20 °C,^5^ in cockroaches at 17 °C,^6^ in *Neurospora* at 10.5 °C,^7^ and in *Arabidopsis*,^8^ the European chestnut *Castanea sativa*,^9^ and tomato *Lycopersicon esculentum Mill* at 4 °C.^10^

Circadian rhythms in homeothermic animals under low-temperature conditions are reported far less frequently. A few studies have addressed mammalian circadian rhythms in the absence of body temperature homeostasis. One notable area of research focuses on circadian rhythms in hibernating rodents. The core body temperature of a hamster, including the brain, approaches 5 °C during hibernation. During this state, circadian rhythms of gene expression within the suprachiasmatic nucleus (SCN) are abolished.^11,12^ More recent studies using calcium live imaging have demonstrated that circadian rhythms in SCN neurons disappear at 15 °C.^13^

The loss of self-sustained oscillation, such as the nullification of circadian rhythm under low temperature conditions, can be regarded as a qualitative change in a dynamical system. Dynamical system theories provide mathematical scenarios that occur due to changes in parameter values, a phenomenon referred to as bifurcation.^14^ Thus, one can expect that bifurcation underlies the arrhythmicity induced by low-temperature conditions. According to bifurcation theory, two typical scenarios can explain the loss of rhythmicity: Hopf bifurcation and saddle-node on an invariant circle (SNIC) bifurcation (Figure 1). In Hopf bifurcation, the oscillation amplitude gradually diminishes until it eventually reaches zero, and a self-sustained oscillator transforms into a damped oscillator. In SNIC bifurcation, oscillations are arrested at a specific phase, where the system transitions from a self-sustained oscillation to an excitable system. During this process, the period extends to infinity, while the amplitude remains almost constant.

**Figure 1.**
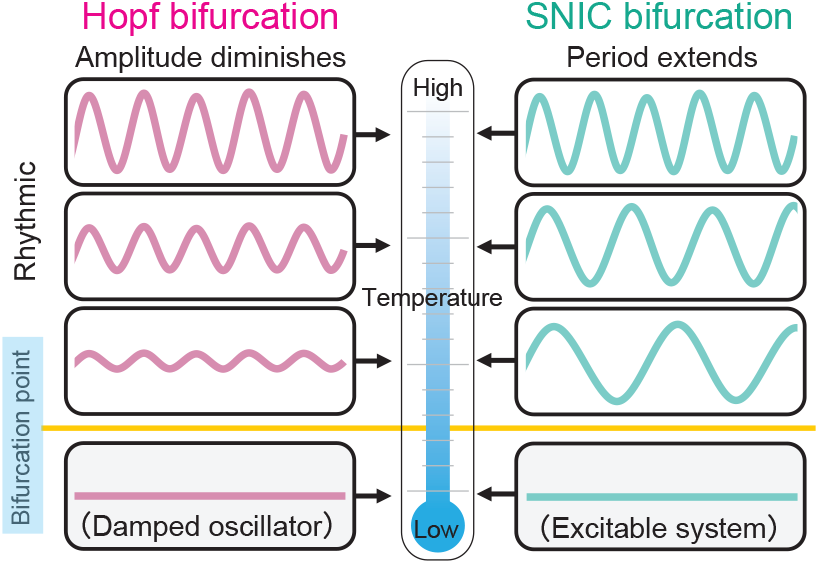
Two potential bifurcations underlying the loss of rhythmicity at low temperatures. The loss of self-sustained circadian rhythms under low temperatures can be explained by two mathematical scenarios. Around the critical temperature, the oscillator behaves differently under the different bifurcations. In Hopf bifurcation, the oscillation amplitude decreases to zero while the period remains relatively stable. In addition, the self-sustained oscillator turns into a damped oscillator at the critical point. Meanwhile, in SNIC bifurcation, the oscillation period extends and eventually diverges to infinity. The self-sustained oscillation can be transformed into an excitable system at the bifurcation point.

A previous study demonstrated that low temperature nullifies the circadian clock of cyanobacteria through Hopf bifurcation,^15^ where the reconstituted KaiC oscillator transitions into a damped oscillator at 19 °C. However, the type of bifurcation underlying qualitative changes in the circadian clock of homeothermic animals has not yet been investigated. In this study, we focused on the loss of circadian rhythmicity in cultured human cells under low-temperature conditions and analyzed the qualitative transitions of the circadian clock using bifurcation theory. Furthermore, we examined human hair follicle cells to investigate the temperature dependence of peripheral circadian clocks in homeothermic organisms under low-temperature conditions. Through these investigations, we explored the universal dynamics governing circadian rhythms under low-temperature conditions and their evolutionary implications.

## RESULTS

### Reduced circadian amplitude of cultured human cells under low temperatures

To determine the bifurcation type involved in the loss of self-sustained oscillations, a detailed analysis of the period and amplitude near the bifurcation point is required. Thus, we cultured U2OS cells expressing the *BMAL1*-dLuc reporter in wells with a controlled temperature gradient and monitored the bioluminescence patterns across a temperature range from 25.1 °C to 36.9 °C (Figure 2A). The oscillatory rhythms became undetectable at approximately 30 °C (Figure 2B). As the temperature decreased, the oscillation amplitude of the bioluminescence rhythm progressively diminished, while the oscillation period did not significantly change. This temperature-dependent reduction in amplitude, coupled with the stability of the period near the point where the rhythm disappeared, suggests that the loss of circadian rhythmicity under low-temperature conditions is governed by Hopf bifurcation.

**Figure 2.**
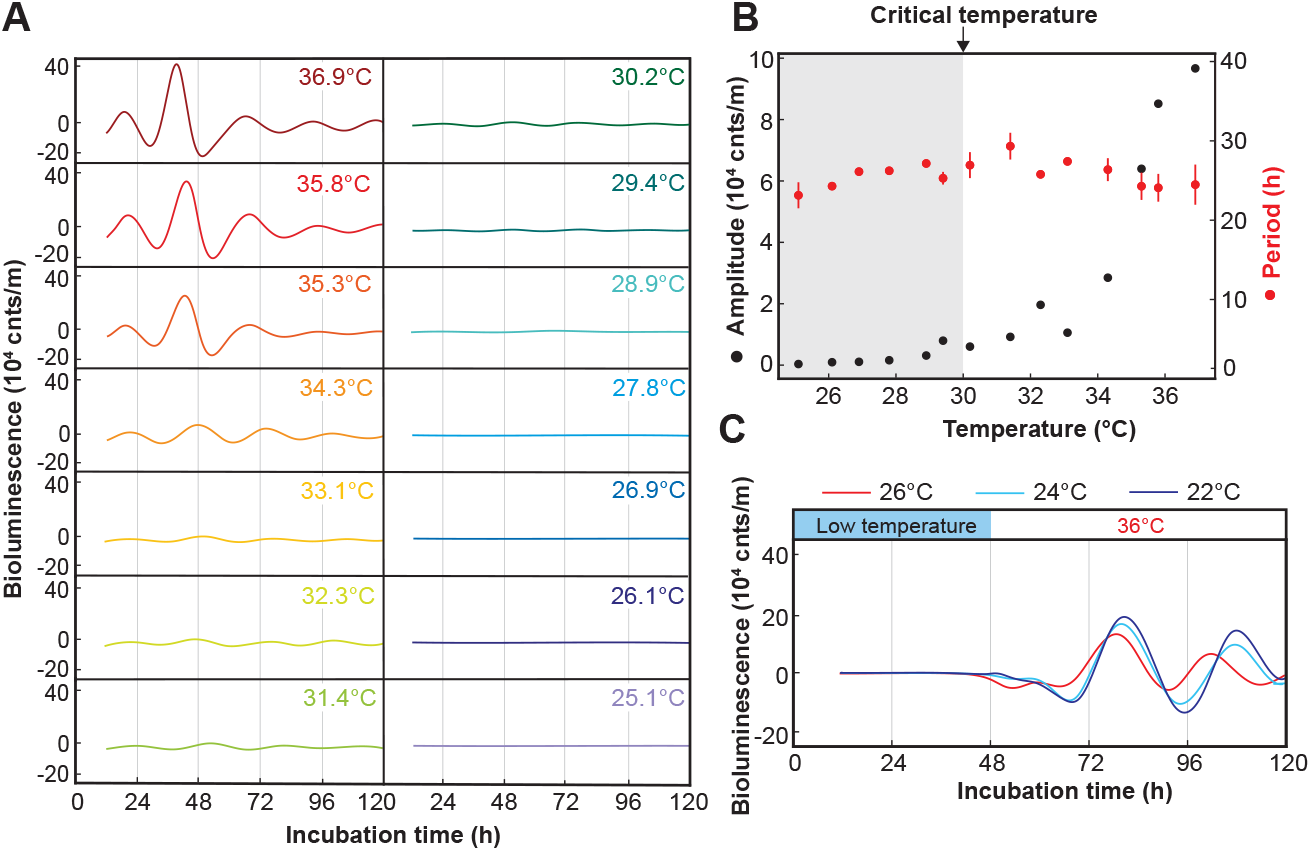
Loss of rhythmicity of cultured human cells under low temperatures. (A) Bioluminescence rhythm of U2OS *BMAL1*-Luc cells at low-temperature conditions. (B) Amplitude (black) and period (red) of the bioluminescence rhythm under different temperatures. The period is presented as the mean ± SD. (C) Recovery from cold-induced arrhythmia. U2OS cells were incubated at 22 °C, 24 °C or 26 C for 48 h and then transferred to 36 °C.

### Recovery from cold-induced arrhythmia with a different phase

When plants or poikilotherms lose circadian rhythmicity under low temperatures and resume the rhythm upon rewarming, the rhythms tend to start from circadian time (CT) 12, the beginning of the subjective night.^4,16^ To investigate whether a similar phenomenon occurs in human cells, we exposed U2OS cells to low temperatures (22–26 °C), followed by a rewarming to 36 °C. Consistent with observations in plants and poikilotherms, rewarming to 36 °C restored the circadian rhythm of *BMAL1* oscillations (Figure 2C).

In addition, we estimated the phase of the *BMAL1* rhythm upon resumption based on its phase after rewarming to 36 °C. Consistent with prior studies,^17,18^ we defined CT22 as the phase value corresponding to the peak of *BMAL1* expression rhythm and estimated the phase at which the rhythm restarted based on CT22 – 24 *t*/*τ*, where *t* represents the time of the first peak after rewarming, and *τ* is the free-running period at 36 °C. Accordingly, we estimated that oscillations began at CTs 12.38, 13.08, and 14.76 for initial temperatures of 22, 24, and 26 °C, respectively. Notably, the phase of the oscillation rhythm after rewarming to 36 °C was not fixed at a specific phase but instead depended on the combination of temperatures. This phenomenon can be attributed to the temperature-dependent position of the stable fixed point created by Hopf bifurcation, rather than by SNIC bifurcation (Figures S1A and S1B).

### Resonance of damped oscillation in response to temperature cycles

According to Hopf bifurcation theory, the bifurcation can transform a self-sustainable oscillator into a damped oscillator at a critical point.^14^ If exposure to low temperature induces Hopf bifurcation in the cellular clock, one can expect resonance of the circadian clock dampened by low temperature, which is analogous to the enhancement of the pendulum’s amplitude by an optimal periodic force. In cyanobacteria, the phosphorylation rhythm of reconstituted KaiC is transformed into a damped oscillator under low temperature conditions, and temperature cycles with an optimal period can restore the amplitude of the damped oscillation.^15^ Similarly, a damped bioluminescence rhythm is observed in *kaiA* deletion mutants of cyanobacteria, which resonate with external temperature cycles with a period of 24-26 h.^19^

Thus, we periodically applied 3-h 37 °C temperature pulses to U2OS chilled at 30 °C, which can be close to the bifurcation point (Figures 3A and S2). The amplitude of the forced oscillation depended on the period of the temperature cycle and peaked at temperature cycles with a period of 24 h (Figure 3B), indicating that the low-temperature-induced damped circadian rhythm exhibited resonance. The resonant frequency should be close to 24 h, which is consistent with the fact that the transition by Hopf bifurcation tends to keep the oscillation period.

**Figure 3.**
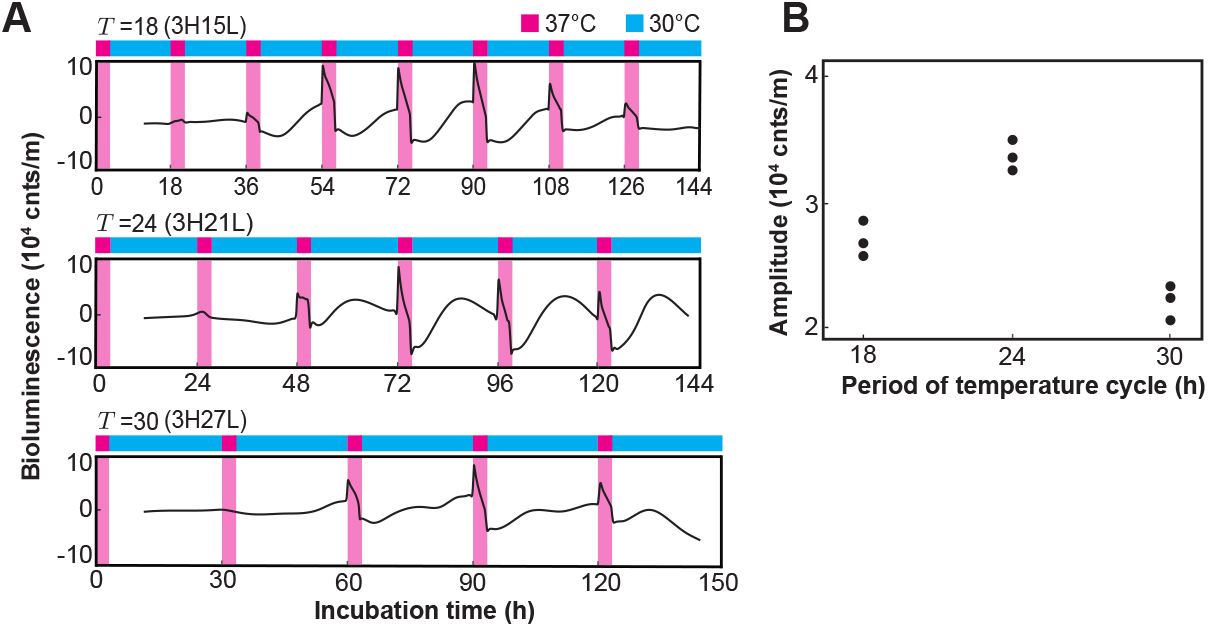
Resonance of abolished bioluminescence rhythm by periodic temperature pulses. (A) Bioluminescence of cultured human cells under periodical 3-h 37 °C pulses with a period of *T* =18, 24, or 30 h (H: 37 °C; L: 30 °C). (B) Amplitude of the forced oscillations under periodical temperature pulses. The amplitude was evaluated by the SD of time course data from 2*T* to 120 h under 30 °C.

### Preference for Hopf bifurcation in the mathematical model

To explore the underlying molecular mechanisms for Hopf bifurcation, we focused on the Kim– Forger model, which describes the dynamics of the detailed molecular network of the mammalian clock.^20^ This comprehensive model incorporates 70 parameters and 180 variables, integrating both positive and negative feedback loops. To investigate the effect of temperature on the dynamics of this model, we assumed that some of the rate constants become smaller as the temperature decreases, i.e., we randomly assigned some parameters in the model as temperature-dependent. The reduction of the parameter value due to low temperature was parameterized by a factor *α*, which ranged between 0 and 1. We assumed that lowering the temperature can update the parameter value according to *k*^*′*^ = *αk*, where *k* and *k*^*′*^ represent the value of the temperature-dependent rate constant under normal and chilling conditions, respectively. We quantified the effect of the reduction of the parameter value on the amplitude and period of *Bmals* mRNA in the nucleus, which was representative of the mammalian circadian clock dynamics. We classified the type of bifurcation based on the transition of oscillation amplitude and period around the critical point of rhythm disappearance (Figures S3 and S4).

Reducing the transcription rate constant of *Bmal* led to a decrease in oscillation amplitude, while the period remained relatively stable at approximately 24 h, suggesting that a Hopf bifurcation occurs around *α* = 0. In contrast, the combination of three specific parameters, namely, reducing the degradation rate constant for BMAL, unbinding the rate constant of REV-ERBs to the *Cry1*/*Npas2* ROR-response element, and unbinding the rate constant of PER1/2 to CRY1/2, caused the oscillation period to diverge to infinity while maintaining a stable oscillation amplitude, suggesting a SNIC bifurcation. The variation in the transcription rate constant of *Per1* did not result in loss of rhythmicity (Figures 4A and S3).

**Figure 4.**
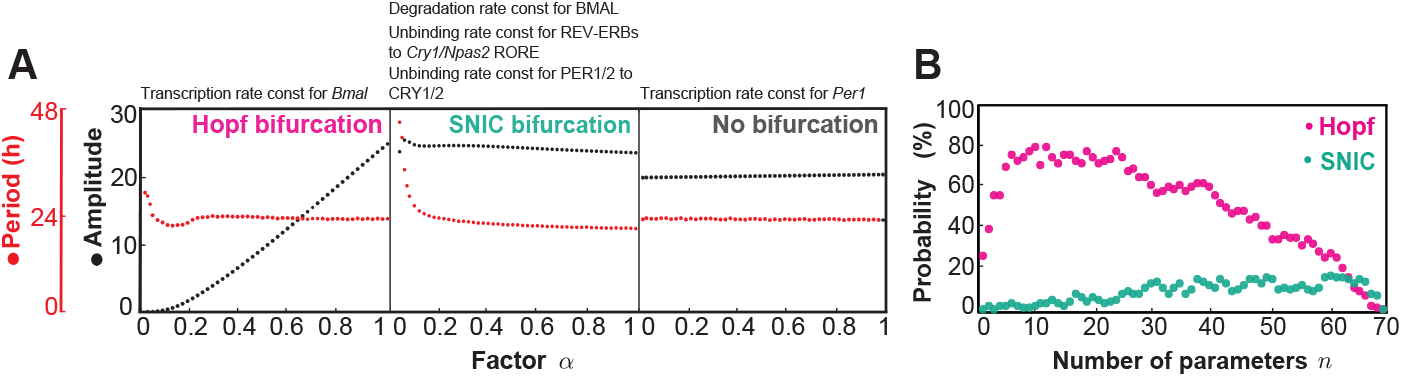
Bifurcation analysis for a detailed mammalian circadian model. (A) Temperature-dependent time course of oscillatory variables for the model proposed by Kim and Forger.^20^ To classify the type of bifurcation, nuclear *Bmals* expression was selected as a representative marker of circadian dynamics. We used the original parameter and hypothesized that some of the 70 parameters of the Kim–Forger model depend on temperature. Here we show the bifurcation examples of transcription rate constant for *Bmal* and *Per1*. In addition, we classified the bifurcation for three parameter sets, degradation of the BMAL rate constant, unbinding rate of PER1/2 to CRY1/2, and unbinding of the rate constant of REV-ERBs to *Cry1*/*Npas2* RORE. (B) Probability of bifurcation type depending on the number of temperature-dependent parameters *n*.

To investigate which type of bifurcation is more likely to occur when reducing parameter values, we randomly selected *n* temperature-dependent parameters 100 times and examined the types of bifurcations that emerged as factor *α* decreased. Hopf bifurcations were frequently observed across a wide range of *n* values. In contrast, SNIC bifurcations occurred with a lower probability compared to Hopf bifurcations (Figure 4B). This preference for Hopf bifurcations was reproduced in the Goodwin model,^21^ which describes a simple transcriptional negative feedback loop (Figure S5). These results support our experimental results and further suggest that biological oscillators based on a negative feedback loop have a tendency to generate Hopf bifurcations.

### Temperature-insensitive circadian amplitude of human hair follicles

While we have investigated the effect of temperature on cultured human cells, the temperature effect on the individual level is more complex. In particular, the temperature dependency of cellular rhythms in the human body remains poorly understood. To address this gap, we investigated circadian rhythms of hair follicle cells in human subjects isolated in a temperature-controlled room. The advantages of hair follicle cells are that the expression level of the follicle cells can be measured non-invasively^22^ and the shallowness of follicle cells in the scalp skin suggests possible dependence on environmental temperature.

As a preliminary experiment, we examined the expression of nine clock genes in hair follicle cells at three time points (Figure S6). We extracted the amplitudes and phases from these time courses and selected four high-amplitude genes. In the subsequent main experiment, we focused on the expression of these selected genes together with the *Cry1* gene, which is known to exhibit temperature-dependent amplitude regulation.^23^ Hair follicle cells were collected every 6 h from six participants who stayed in temperature-controlled chamber maintained at either 28 °C or 18 °C (Figure 5A). The scalp skin temperature showed a circadian rhythm, and its average level depended on the ambient temperature, as expected (Figure 5B). Notably, the timing of the trough of scalp temperature rhythm was delayed under cooler conditions, and the thermal fluctuations were larger at 18 °C than those at 28 °C.

**Figure 5.**
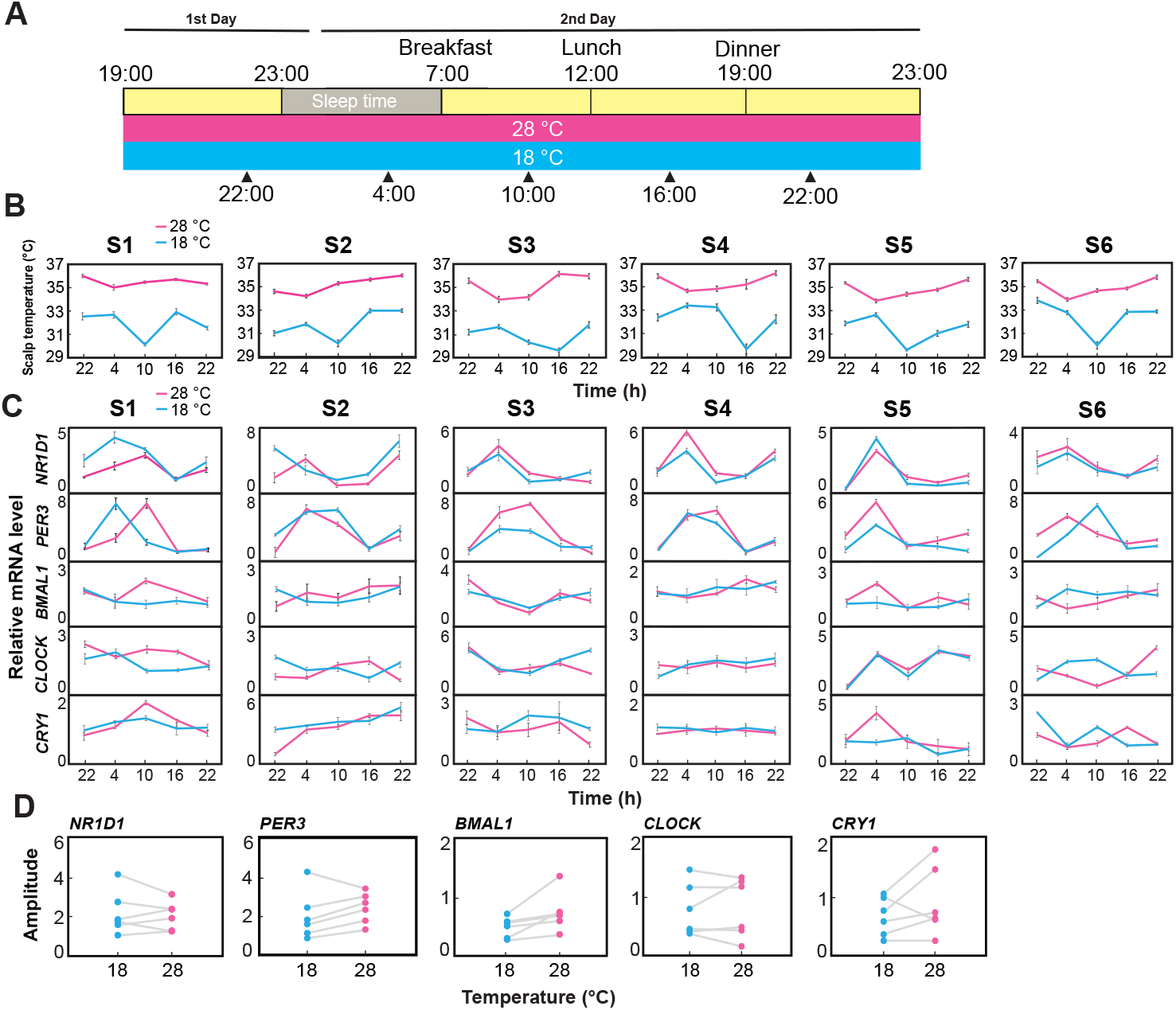
Circadian rhythms of hair follicles in a temperature-controlled chamber. (A) Hair follicle collection procedure. Six participants arrived at the experimental facility at 19:00 and were dressed in experimental clothes after receiving brief instructions for the experiment. The participants stayed in the isolated temperature-controlled chamber (28 °C and 18 °C) until the end of the experiment. Ten hair follicles were collected from participants at 22:00, 4:00, 10:00, 16:00, and 22:00. The hair follicles samples were immediately soaked in RNAlater and stored at -70 °C until RNA analysis. (B) Scalp skin temperature under different temperatures. Each participant’s scalp skin temperature was measured three times at one-minute intervals. The value is presented as the mean ± SD. (C) Expression profiles of the circadian clock gene under different temperatures. The error bars indicate the SD of three repeated measurements by real-time qPCR. (D) Circadian amplitude of gene expression profiles under different temperatures. Each dot represents the circadian amplitude of hair follicle cells from an individual participant, shown as the SD. Gray lines connect the amplitudes of the same participant under different temperature conditions.

We quantified mRNA expression levels of key circadian regulators (*NR1D1, PER3, BMAL1, CLOCK*, and *CRY1*) in human hair follicles under 28 °C and 18 °C (Figure 5C). The amplitude of these expression profiles revealed no statistical significant differences between different temperature conditions (Figure 5D), demonstrating that the circadian amplitude of hair follicles in the human body maintained thermal stability.

We further investigated the circadian amplitude of ex vivo cultured human hair follicles under different temperature conditions (Figure S7A). The amplitudes of the expression profile of *PER3* and *CLOCK* were enhanced at 36 °C compared with that of 26 °C. Conversely, the amplitudes of *BMAL1* and *NR1D1* were smaller at 36 °C than at 26 °C (Figure S7B), suggesting different temperature dependency in the amplitude compared to cultured U2OS cells. These observations imply that the circadian amplitude of ex vivo cultured human hair follicles is more sensitive to ambient temperature changes than those in in vivo conditions, and they exhibit different sensitivities between clock genes.

## DISCUSSION

We demonstrated that low temperature diminished the oscillation amplitude of circadian rhythms in cultured human cells, suggesting that Hopf bifurcation underlies the loss of rhythmicity. In addition, the resonance of the 24-h temperature cycle resulting in the enhancement of the circadian amplitude supports the involvement of Hopf bifurcation. Considering the phosphorylation rhythm of cyanobacterial KaiC,^15^ these observations imply that two distinct clock mechanisms exhibited low-temperature-induced Hopf bifurcation. In Hopf bifurcation, the oscillation period remains relatively constant near the bifurcation point, whereas significant period slowing is observed in SNIC bifurcation. Given that compensation across a wide range of temperatures is a critical adaptive feature of circadian rhythms, it is plausible that the circadian system prefers Hopf bifurcation. These observations suggest Hopf bifurcation is a universal property along with temperature compensation and entrainment to environmental cycles.

The Kim–Forger model confirmed the preference for Hopf bifurcation (Figure 4B). The vulnerability of amplitude and robustness of period in this model can be observed even far from the bifurcation point,^23^ which suggest the association between temperature compensation and Hopf bifurcation. Moreover, because the Kim–Forger model comprises multiple negative feedback loops, temperature compensation and Hopf bifurcations may naturally arise as inherent properties associated with such multiple feedback loops.

While the qualitative loss of rhythmicity can be conserved across species, the critical temperature is not qualitatively conserved. For example, we showed that the critical temperature of U2OS cells was approximately 30 °C (Figure 2B). However, a previous study reported that the critical temperature of SCN neuron slices was 15 °C.^13^ The circadian rhythm in hibernating ground squirrels and bats continues to function at body temperatures around 10 °C.^24,25^ Hibernating European hamsters do not show circadian oscillations when the core body temperature drops to near-ambient temperature (5 °C).^12^ These observations suggest that the critical temperature depends on levels of the organization, i.e., cellular, tissue, or organismal level. Such diversity in critical temperatures may stem from the variability in the amplitude of circadian clocks. A larger amplitude would likely result in a lower critical temperature. Moreover, it is known that some clock genes are involved in determining the amplitude of the circadian clock.^26–28^

Resonance to temperature cycles with an optimal frequency enhanced the circadian amplitude of cultured human cells (Figures 3A and 3B). Resonance can transfer energy between oscillators, which provides an energetic advantage for biochemical oscillations.^29,30^ Therefore, one can expect physiological merits for resonating with environmental frequency. Plants and cyanobacteria under light-dark cycles with periods close to the endogenous circadian rhythm exhibit enhanced growth.^31–33^ Conversely, deviations to the innate 24-h circadian period in non-human primates and rodents resulted in reduced lifespans.^34–36^ These connections between the environmental period and physiological advantage can be explained by the enhancement of amplitude by resonance. Winter coldness can severely reduce circadian amplitude in plants or poikilotherms.^4,15,37^ However, resonance could recover this reduction of oscillation amplitude. Moreover, resonance to temperature cycles may function in homeothermic animals. The circadian clock in mammals rhythmically regulates body temperature. The resulting body temperature rhythm, in turn, is expected to enhance the circadian clock’s amplitude through resonance. This interaction between the circadian clock and body temperature rhythm via resonance may be one of the evolutionary reasons why peripheral clocks in homeothermic animals exhibit temperature sensitivity.

From a more microscopic view, the causes of amplitude reduction near a Hopf bifurcation are generally classified into two cases: amplitude reduction of individual oscillators or desynchronization, i.e., loss of rhythmicity in cultured cells caused by desynchronization between cells, or decrease in amplitude in each cell. Decreased oscillation amplitude in individual SCN neurons at low temperature suggests oscillation death in each cell.^13^ However, further single-cell observation around critical temperature is needed to elucidate the cause of Hopf bifurcation.^38^

We demonstrated that an ambient temperature of 18 °C reduced scalp skin temperature yet did not affect the amplitude of clock gene expression in hair follicle cells (Figure 5). This suggests that the follicle cells, located in the dermal layer of the skin and influenced by blood flow,^39^ can be maintained under relatively stable temperature conditions regardless of ambient temperature. The actual temperature measurements on or within the scalp are consistent with this suggestion.^40^ In contrast, the circadian amplitude in ex vivo cultured hair follicle cells was more sensitive to external temperature fluctuations (Figure S7B). This suggests that isolated cells lack the integrative mechanisms required to buffer against environmental temperature changes. Temperature homeostasis in humans should contribute to maintaining the amplitude of peripheral clocks in the whole body by resisting the temperature dependence of circadian amplitude.

### Limitations of the study

Bioluminescence assays driven by luciferase reporter genes were utilized as a standard approach to monitor circadian rhythms.^41,42^ However, because bioluminescence depends on metabolic activity and is temperature-sensitive, it may confound amplitude measurements and bifurcation analysis. Developing temperature-independent methods is crucial for accurately characterizing core circadian oscillator dynamics and defining bifurcation types.

Although our experiments using U2OS cells and hair follicle cells provided valuable insights into temperature-dependent circadian regulation, these models do not fully replicate the complexity of whole-body systems or the behavior of tissues with distinct thermal sensitivities, potentially limiting the generalizability of the findings.

While the study confirmed robust temperature homeostasis in human hair follicle cells in vivo, the precise temperature within the human hair follicles during environmental fluctuations was not directly measured, which limits accurate correlation between ambient temperature and in vivo circadian dynamics.

## STAR METHODS

### Key resources table

| Reagent or Resource | Source | Identifier |
| --- | --- | --- |
| **Cell Lines** |  |  |
| Human U2OS cells | ATCC | HTB-96 |
| Human hair follicle cells | Primary culture | N/A |
| **Reagents** |  |  |
| Luciferin | Thermo Fisher Scientific | L2916 |
| RNAlater stabilization solution | Thermo Fisher Scientific | AM7020 |
| RNAqueous Micro kit | Thermo Fisher Scientific | AM1931 |
| Superscript IV VILO Master Mix | Thermo Fisher Scientific | 11756050 |
| TB Green Premix EX Taq II | TaKaRa | RR820L |
| Dulbecco's Modified Eagle Medium (DMEM) | Thermo Fisher Scientific | 3090869 |
| Fetal bovine serum (FBS) | Biosera | FB-1290/500 |
| **Equipment** |  |  |
| Photomultiplier tube | Hamamatsu | H11901-01 |
| CO<sub>2</sub> incubator | Thermo Fisher Scientific | Heracell 150i |
| Bio Rad thermocycler | Thermo Fisher Scientific | C1000 Thermal cycler |
| Heating baths | Ampere | UT40U100F |
| **Software and Algorithms** |  |  |
| MATLAB | MathWorks | R2024b |
| GraphPad Prism | GraphPad | Version 9 |

## RESOURCE AVAILABILITY

### Lead contact

Requests for further information and resources should be directed to and will be fulfilled by the lead contact, Hiroshi Ito.

### Materials availability

This study did not generate new unique reagents.

### Data and code availability

The code reported in this paper is available from the lead contact upon request or directly at: https://github.com/hitolab/humanclock. Any additional information required to reanalyze the data reported in this paper is available from the lead contact upon request.

### Experimental model and study participant details

#### Collection of human hair follicle cells from the scalp

Six healthy participants (three male and three female university students) were chosen for this experiment. The participants ranged in age from 22 to 32 years. The Kyushu University Institutional Review Board for Human Genome/Gene Research approved this study protocol (approval number: 548-08), and all procedures were carried out in accordance with the approved guidelines. After describing the experimental procedure to experimental participants, written informed consent was obtained from all subjects prior to enrolment.^43^

All participants stayed in a homotoro chamber (temperature-controlled room) for one day. The room was set to a high temperature of 28 °C or a low temperature of 18 °C. A temperature detector was used to measure their scalp skin temperature. Before the experiment, all participants were asked to follow a sleep-wake schedule between 23:00 and 07:00 for at least one week. Each participant wore an Actiwatch Fitbit Charge 5 during the study period. An actigraph was used to confirm whether they followed the schedule. The participants were not allowed to consume excessive alcohol, snacks, or caffeine before the experiment. Ten scalp hair follicles were collected from each subject by holding and pulling hair with a pair of tweezers. The hair shafts largely covered with follicle cells were quickly immersed in RNAlater™ Stabilization Solution (AM7020, Thermo Fisher Scientific, U.S.A) and stored below -20 °C until RNA purification.

### Cell culture of U2OS carrying *BMAL1* luciferase promoter

Human U2OS cells were provided by Dr. Takashi Yoshimura.^44^ The cells were cultured in Dulbecco’s Modified Eagle Medium (DMEM; 3090869, Thermo Fisher Scientific, USA) supplemented with 10% fetal bovine serum (FBS, FB-1290/500; Biosera, France), 100 U/mL penicillin, and 100 µg/mL streptomycin (Pen Strep, 15070-063, Thermo Fisher Scientific, Waltham, MA, USA) for several days at 37 °C incubator with 5% CO_2_.

### Method details

#### Real-time monitoring of *BMAL1*-Luc oscillations in U2OS cells

When U2OS cells reached confluence, they were further incubated in serum-free DMEM for 24 h and then synchronized using 100 nM dexamethasone (Dex, Sigma-Aldrich). After 2 h of treatment with Dex, the culture medium was replaced with complete DMEM containing 10% FBS and 1% penicillin-streptomycin solution, and the dish was placed in a photomultiplier tube (PMT) under different temperatures. Luciferase activity was monitored using a real-time PMT. The data are presented as counts per minute, calculated at 10-minute intervals. A baseline correction was calculated using a 24-h moving average, which removed the first 12 h of data, as previously described.^45^

#### Measurement of amplitude and period for bioluminescence

The amplitude was examined by standard deviation (SD), and the period was determined using peak-to-peak fluctuations. We calculated the SD of all detrended data of bioluminescence as circadian amplitude. The period was determined based on the difference between peaks using the scipy.signal.find peaks function in the Python3 module.

#### Temperature-gradient apparatus

Two heating baths (UT40U100F; Ampere) were connected by an aluminum block insulated with a heat-resistant material.^15^ By adjusting the temperature of the heating baths, a precise temperature gradient was established with an accuracy of ±0.1 °C.

#### Isolation of total RNA, reverse transcription PCR, and real-time quantitative PCR

Total RNA was extracted from human hair follicles using the RNAqueous-Micro Kit (Thermo Fisher Scientific, U.S.A). The cDNA sample was generated by using SuperScript™ IV VILO™ Master Mix. The primer sets used for real-time quantitative PCR are based on a previous study.^46^ Briefly, qPCR was performed in a 20 µL reaction volume containing TB Green premix Ex Taq II (Tli RNaseH Plus), primers, template, and sterile purified water using the CFX96 Real-time PCR Detection System (Bio-Rad), according to a previously described method.^45^ All reactions were performed in triplicate and displayed amplification efficiencies between 80% and 120%. The 2^*−*ΔΔ*Ct*^ method was used to quantify gene expression. In addition, *GAPDH* was used as an internal reference to normalize the relative expression of each sample.

#### Numerical analysis of the mammalian clock model

Kim–Forger model was numerically solved using the MATLAB code provided by the authors.^20^ To determine the type of bifurcation, we chose the concentration of *Bmals* in the nucleus as a representative variable of circadian dynamics. We hypothesized that some of 70 parameters of the Kim–Forger model depend on temperature, i.e., *k*^*′*^ = *αk* where *k* represents the value of temperature-dependent parameter and *α* ∈[0, 1] is a temperature-dependent factor, with 1 corresponding to high temperature and 0 to low temperature. We adopted the value of the rate constants described in the original paper.^20^ In addition, we calculated the probability of bifurcation by randomly selecting temperature-dependent parameters (Figure 4B). We determined the type of bifurcation for each set of temperature-dependent parameters through the procedure shown in Figure S4.

We considered Goodwin model with three variables represented as

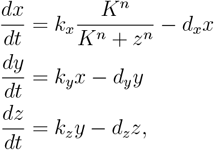

where *x* represents the amount of clock gene mRNA, *y* and *z* represent the amount of clock proteins in the cytoplasm and nucleus, respectively. *k*_*∗*_ and *d*_*∗*_ are the rate constants for synthesis and degradation, respectively. *K* is the apparent dissociation constant and *n* is the hill coefficient. We set original parameter values as follows: *k*_*x*_ = *k*_*y*_ = *k*_*z*_ = *K* = 1, *d*_*x*_ = *d*_*y*_ = *d*_*z*_ = 0.1, *n* = 10. We assumed that some of the rate constants, *k*_*x*_, *k*_*y*_, *k*_*z*_, *d*_*x*_, *d*_*y*_, *d*_*z*_ in Goodwin model were temperautre-sensitive in the same manner as Kim-Forger model, i.e., *k*^*′*∗^ = *αk*_*∗*_.The solver in MATLAB, `ode15s`, was used for numerical simulations for Goodwin model.

The `findpeaks` function in MATLAB was used to detect the peaks and measure the oscillation period from the numerical simulations of both models.

### Quantification and statistical analysis

All the statistical analyses were performed using Excel (Microsoft) and GraphPad Prism 9 (GraphPad Software). The values are presented as the mean ± SD. Statistical details of experiments can be found in the figure legends, figures, and the results section.

## ACKNOWLEDGMENTS

The authors thank T.N. Ohkawa, T. Yoshimura, and M. Akashi for providing U2OS cells and the human *PER3* plasmid. This work was supported by the Japan Society for the Promotion of Science (JSPS) KAKENHI (grant JP23H04475) to H.I., AMED CREST JP24gm2010005 to H.I., and a scholarship from the Chinese Government Scholarship Council (grant number: 202006300020) to Y.X.

## AUTHOR CONTRIBUTIONS

Conceptualization, Y.X. and H.I.; methodology, Y.X., T.N., and H.I.; investigation, Y.X., Y.S., and H.I.; writing–original draft, Y.X. and H.I.; writing–review & editing, Y.X. and H.I.; funding acquisition, H.I.; supervision, H.I.

## SUPPLEMENTAL INFORMATION INDEX

Figures S1–S6 and their legends in a PDF

