## Supplemental information for "Low temperature abolishes human cellular circadian rhythm through Hopf bifurcation"

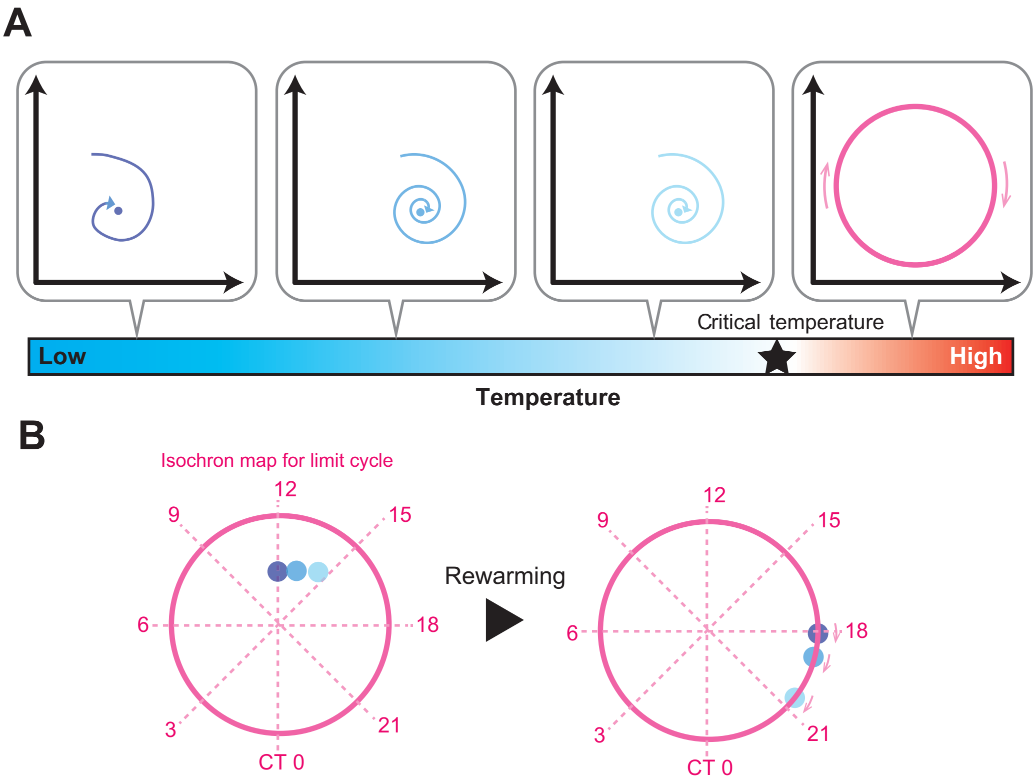


**Figure S1. Cellular rhythm dynamics during rewarming. Related to Figure 2.** (A) According to Hopf bifurcation, a limit cycle oscillator turns into a damped oscillator at a critical point. Thus, we can expect that the circadian oscillator is a damped oscillator below the critical temperature. The trajectory of self-sustained oscillation can be represented by a closed orbit on a phase plane, whereas damped oscillation can be a spiral that approaches a stable fixed point at $t \to \infty$. We assume that the location of the generated fixed points depends on the pre-incubation temperature. (B) A limit cycle and the stable fixed points before and after rewarming. The dotted lines represent hypothetical isochrons for the limit cycle. The result shown in Figure 2C suggests that before rewarming, the circadian system stayed at stable fixed points located at CT 12.38, CT 13.08, and CT 14.76. The difference in the location of fixed points can lead to variations in the phase of the recovered rhythm.

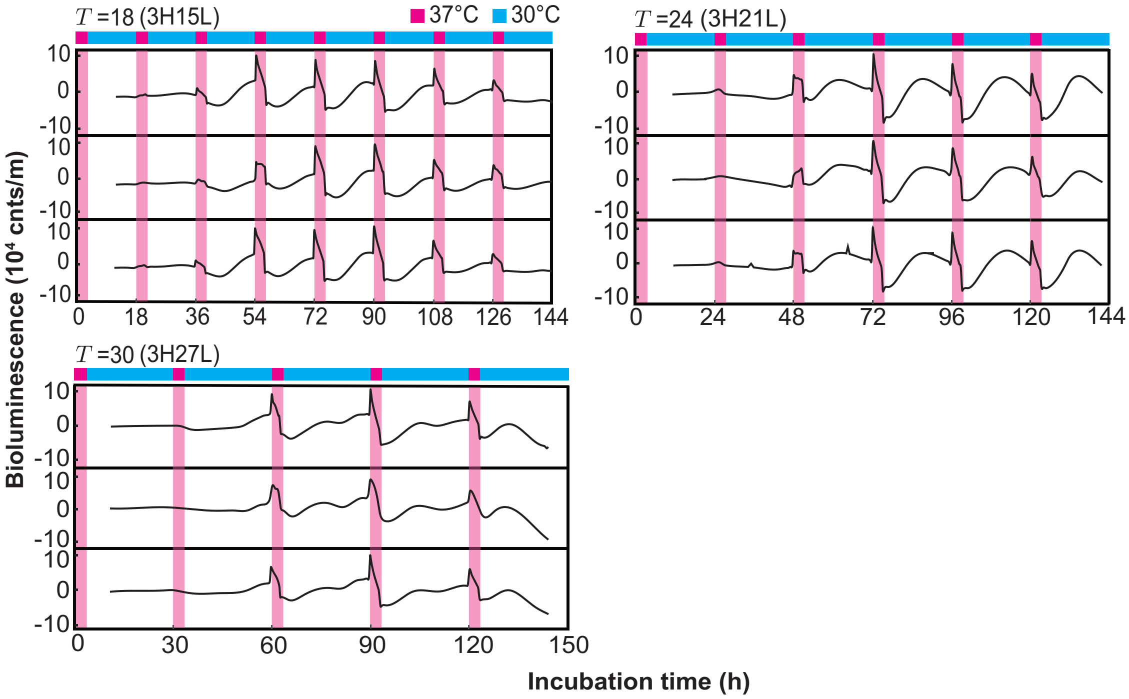


**Figure S2. Resonance of damped oscillation in response to periodic temperature pulses. Related to Figure 3.** Bioluminescence was monitored under periodic 3-h 37 °C pulses with a period of $T$ = 18 h, 24 h, and 30 h.

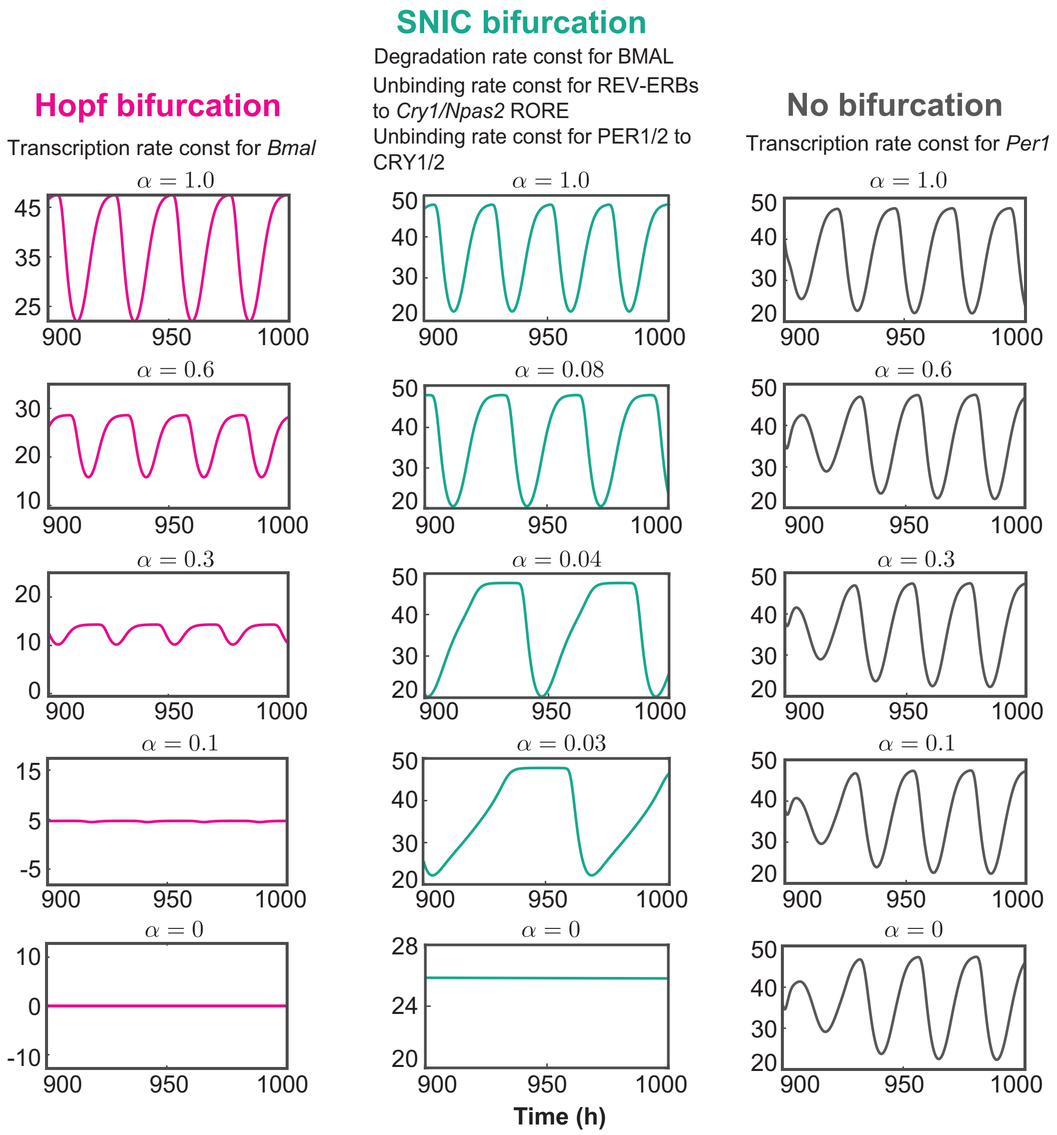


**Figure S3. Examples of bifurcation in the Kim–Forger model. Related to Figure 4A.** We chose *Bmals* mRNA expression in the nucleus as an output variable to investigate the type of bifurcation. The dynamics depended on the parameter value of the temperature-dependent constant rate, $k' = \alpha k$ where $\alpha \in [0, 1]$ is a temperature-dependent factor and $k$ is an original rate constant at normal temperature.

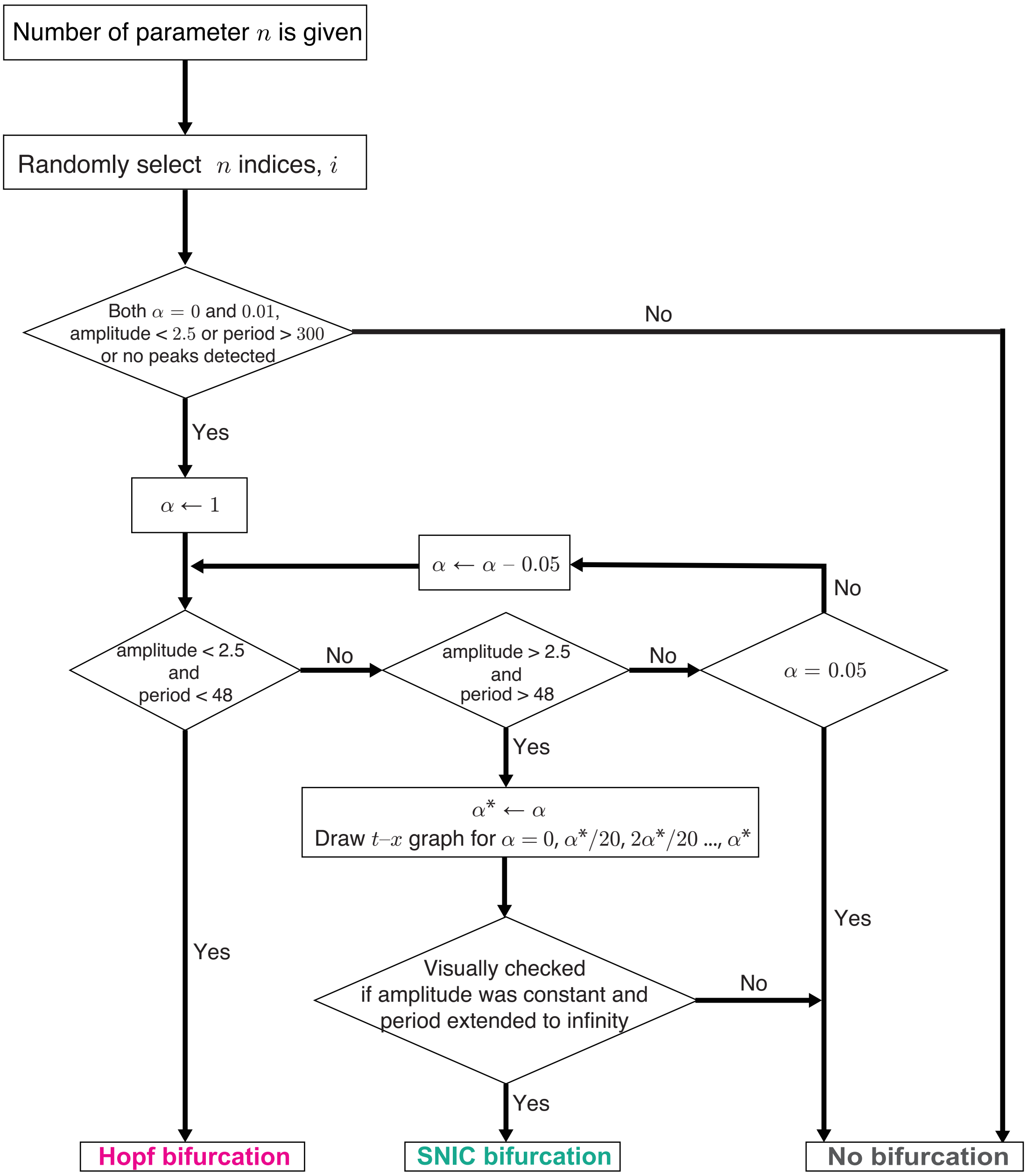


**Figure S4. Flowchart for classifying the bifurcation type in the numerical simulation. Related to Figure 4.** We focused on the nuclear *Bmals* mRNA expression during $900 \leq t \leq 1000$ to avoid the transient period. The peak-to-peak intervals were measured to detect the oscillation periods. Oscillation peaks were detected using the `findpeaks` function in Matlab. The oscillation amplitude was defined as the difference between the maximum and minimum value of *Bmals* during the time course.

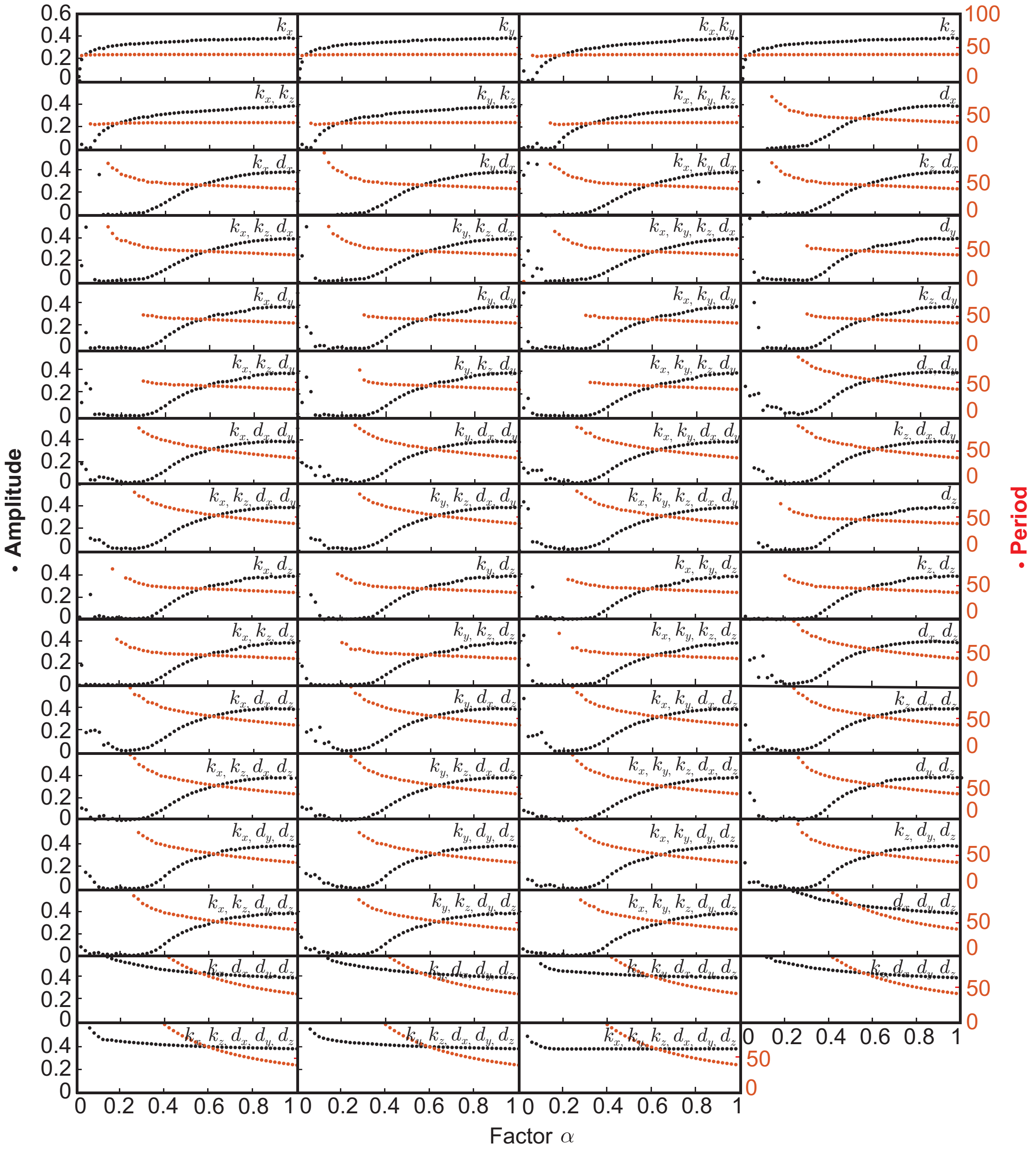


**Figure S5. Bifurcation diagram for Goodwin model.** We selected some rate constants from $k_x, k_y, k_z, d_x, d_y, d_z$ in Goodwin model and altered these parameter values according to $k'_* = \alpha k_*$. The oscillation period and amplitude were observed for different values of $\alpha$. The controlled parameters are shown in each graph.

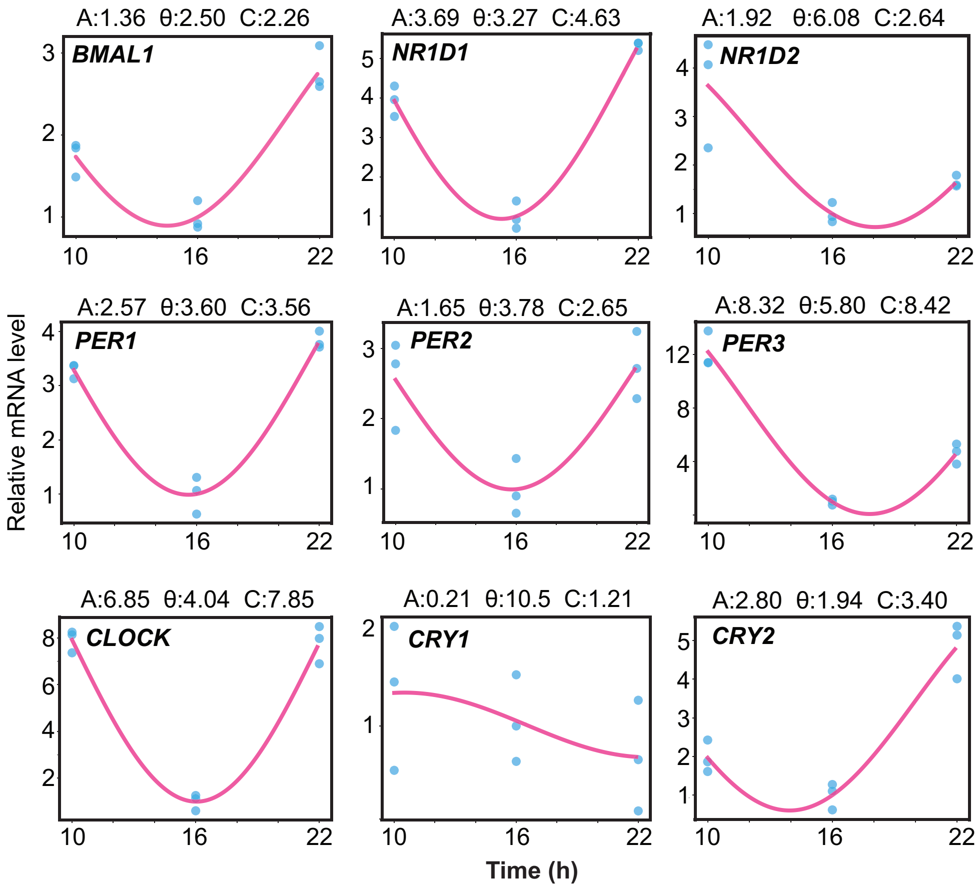


**Figure S6. Expression of circadian clock genes in hair follicles at three time points. Related to Figure 5.** The relative mRNA expression level (blue dots) of the clock genes (*BMAL1*, *CLOCK*, *NR1D1*, *NR1D2*, *PER1*, *PER2*, *PER3*, *CRY1*, and *CRY2*) was normalized to the housekeeping gene *GAPDH*. We fit $A\cos(2\pi(t+\theta)/24)+C$ to determine the amplitude and phase of the oscillations.

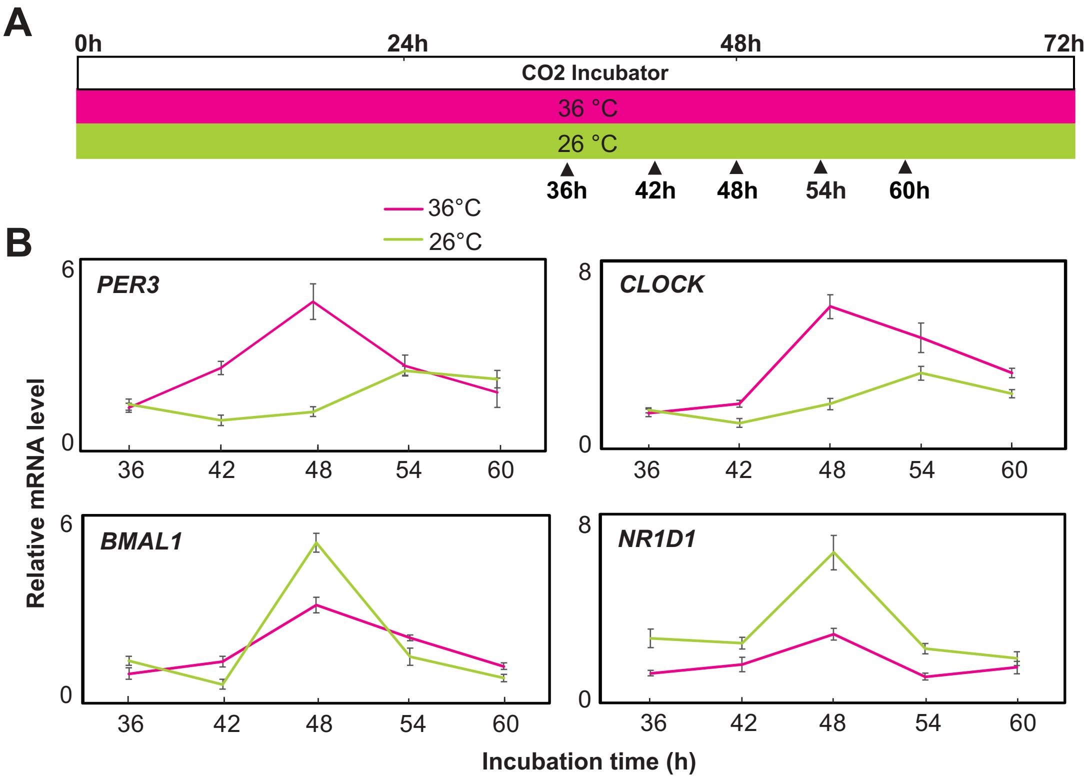


**Figure S7. Circadian rhythms in ex vivo cultured hair follicles under different temperatures. Related to Figure 5.** (A) Sampling schedule of cultured hair follicles. We cultured the hair follicles at 36 °C or 26 °C, and sampled at incubation times of 36, 42, 48, 54, and 60 h. (B) Expression profiles of the circadian clock genes (*PER3*, *CLOCK*, *BMAL1* and *NR1D1*) in the cultured hair follicles.
